# Learning Impairments in *Fmr1^-/-^* mice on an Audio-Visual Temporal Pattern Discrimination Task

**DOI:** 10.1101/2024.09.25.615092

**Authors:** William Mol, Sam Post, Megan Lee, Anubhuti Goel

## Abstract

Estimating time and making predictions is integral to our experience of the world. Given the importance of timing to most behaviors, disruptions in temporal processing and timed performance are reported in a number of neuropsychiatric disorders such as Schizophrenia, Autism Spectrum Disorder (ASD), Fragile X Syndrome (FXS), and Attention-deficit Hyperactivity Disorder (ADHD). Symptoms that implicitly include disruption in timing are atypical turn-taking during social interactions, unusual verbal intonations, poor reading, speech and language skills, inattention, delays in learning, and difficulties making predictions. Currently, there are no viable treatments for these symptoms, the reason being the underlying neural dysfunction that contributes to timing deficits in neuropsychiatric disorders is unknown. To address this unknown, we have designed a novel Temporal Pattern Discrimination Task (TPSD) for awake-behaving mice. Stimuli consist of audiovisual stimuli that differ in duration. Compared to Wild-Type (WT) mice, *Fmr1^-/-^* mice, a well-established mouse model of FXS, showed significant impairment in learning the TPSD task, as evidenced by reduced discriminability indices and atypical licking patterns. Often sensory information is multimodal and indeed studies show that learning in humans and rodents improves with multimodal stimuli than with unimodal stimuli. To test how the multimodal nature of stimuli impacted performance of *Fmr1^-/-^* mice, following training on the audiovisual stimuli, we tested mice on audio-only or visual-only stimuli. While WT mice showed significant disruption in performance when tested on unimodal stimuli, *Fmr1^-/-^* mice displayed equivalent performance on visual-only stimuli when compared to the multimodal task. Our novel task captures timing difficulties and multisensory integration issues in *Fmr1^-/-^* mice and provides an assay to examine the associated neural dysfunction.

## Introduction

Keeping track of the time or duration of events is crucial in predicting an upcoming event. For example, as we wait at a traffic signal our experience includes processing the color of the traffic light (is it yellow or green) but also its duration (how long was the yellow light on for or what was the duration of the stimulus), in order to decide whether to continue pressing the gas pedal or to press the brake. Another example that underscores the importance of time is speech comprehension which relies on learning sequences of syllables that are highly temporally structured–the timing between syllables and duration of each syllable is key to understanding speech. Therefore, an important hallmark of learning is being sensitive to and remembering the temporal structure of events, so that we can make expectations and guide our future decisions. As a result, it’s not surprising that disruptions in timing and timed performance are associated with a number of neurological disorders such as Parkinson’s, Schizophrenia, and Autism. Specifically, disorders of speech perception in Autism are linked to temporal processing deficits (Allman and DeLeon, 2009; Allman et al., 2011; Allman and Meck, 2012). In the context of Attention deficit Hyperactivity Disorder (ADHD), Fragile X Syndrome (FXS), and Autism Spectrum Disorder (ASD), impaired estimation of stimulus duration contributes to atypical perceptual timing, motor timing, and temporal foresight and this is observed on time scales ranging from milliseconds, seconds and days (Hinault et al., 2023). The resulting symptoms include atypical turn-taking during social interactions, unusual verbal intonations, poor reading and language skills, inattention, delays in learning and making predictions, and an inability to incorporate changes to a timed schedule (Sullivan et al., 2006; Puyjarinet et al., 2017; Hoffmann, 2022). While timing tasks demonstrate timing impairments in FXS, ASD, and ADHD, there is a huge gap in our scientific understanding of how dysfunctional neural communication disrupts the representation of time and temporal structure or sequence of events, particularly in the subsecond range. To fill this gap, we have developed a novel rodent timing task, the Temporal Pattern Sensory Discrimination (TPSD) task, where mice have to discriminate between temporal durations of stimuli in order to achieve expert task performance. We found that temporal information was encoded in distinct patterns or trajectories of neural activity, that emerged as mice learned to perform the task (Post et al., 2023). Here we implement the TPSD task to test timing difficulties in FXS, using the well-established model–the *Fragile X* Messenger Ribonucleoprotein 1 *gene* (*Fmr1)* knockout (KO) mouse (Dutch-Belgian Fragile X Consortium, 1994). This model is popular for many reasons. The mouse *Fmr1* gene product shares 97% homology with human FMRP including a conservation of the CGG repeats(Ashley et al., 1993). *Fmr1*-/- mice show functional alterations that are similar to humans and hypersensitivity and few perseverative phenotypes in mice resonate with human symptoms (Dutch-Belgian Fragile X Consortium, 1994; Kazdoba et al., 2014; Goel, 2023). Animal models like mice allow for more granular recording methods and a broader and more nuanced toolkit of experimental manipulations.

The sensory world as it is experienced is rarely restricted to one modality and requires the integration of numerous streams of information and multiple sensory modalities. Persons with ASD have categorically demonstrated reduced capacity for multisensory integration (MSI), symptomatic of and contributory to altered sensory processing generally, across several studies. Many of these studies utilize well-characterized audio-visual illusions such as the Mcgurk effect (Woynaroski et al., 2013; Stevenson et al., 2014b)and sound-induced flash illusion (SiFi) (Foss-Feig et al., 2010; Stevenson et al., 2014a) showing reduced susceptibility to these phenomena, indicative of failure in MSI present in typically developing individuals. Interestingly, these examinations are rich with evidence of atypical receptive temporal processing symptoms, with a marked correlation being established between an expanded temporal binding window and degrees of susceptibility to a particular illusion in individuals with ASD (Foss-Feig et al., 2010; de Boer-Schellekens et al., 2013; Stevenson et al., 2014a). Temporal processing deficits paired with irregularities in audio-visual integration, especially at crucial developmental time points, could provide the basis for impaired linguistic perception and speech reception, both very common in individuals with ASD. Our data shows that similar to psychophysical studies in humans, *Fmr1*^-/-^ mice do not show behavioral facilitation when presented with audio-visual stimuli and rely on visual modality alone for task performance. This assay captures important features of timing and multisensory integration impairment in *Fmr1*^-/-^ mice and provides the framework for future studies to examine the underlying neural dysfunction.

## Methods

### Experimental Animals

All experiments followed the U.S. National Institutes of Health guidelines for animal research, under animal use protocols approved by the Chancellor’s Animal Research Committee and Office for Animal Research Oversight at the University of California, Riverside (ARC #95, respectively). Experiments used male and female FVB.129P2 WT mice (JAX line 004828) and *Fmr1^-/-^* mice (Dutch-Belgian Fragile X Consortium, 1994) (JAX line 004624). All mice were housed in a vivarium with a 12/12 h light/dark cycle, and experiments were performed during the light cycle. The FVB background was chosen because of its robust breeding, because FVB *Fmr1^-/-^* dams are less prone to cannibalizing their pups, and because FVB *Fmr1^-/-^*mice have well-documented deficits in sensory processing (Contractor et al., 2015). We used separate homozygous litters of WT and *Fmr1^-/-^* mice rather than littermate controls because littermates of different genotypes tend to receive unequal attention from the dam (Zupan and Toth, 2008), which may affect the health and behavior of *Fmr1^-/-^* pups, biasing results. To avoid issues with genetic drift, we obtained new WT and *Fmr1^-/-^*breeders from Jackson Labs at regular intervals (every 1-1.5 years).

### Go/No-go temporal pattern sensory discrimination (TPSD) task for head-restrained mice

Awake, head-restrained young adult mice (2-4 months) were allowed to run on an air-suspended polystyrene ball while performing the task in our custom-built rig (**Fig. 1A**). Prior to performing the task, the animals were subjected to handling, habituation, and pretrial phases (Goel et al., 2018; Post et al., 2023; Rahmatullah et al., 2023). After recovery from headbar/cranial window surgery, mice were handled gently for 5 min every day, until they were comfortable with the experimenter and would willingly transfer from one hand to the other to eat sunflower seeds. This was followed by water deprivation (giving mice a rationed supply of water once per day) and habituation to the behavior rig. During habituation, mice were head-restrained and acclimated to the enclosed sound-proof chamber and allowed to run freely on the 8 cm polystyrene ball. Eventually, mice were introduced to the lickport that dispensed water (3-4 µL) and recorded licking (custom-built at the UCLA electronics shop), followed by the audio-visual stimuli. This was repeated for 10 min per session for 3 days. Starting water deprivation prior to pretrials motivated the mice to lick (Guo et al., 2014). After habituation and ∼15% weight loss, mice started the pretrial phase of the training. During pretrials, mice were shown the preferred stimulus only with no punishment time associated with incorrect responses. This was done in order to 1) teach the mice the task structure and 2) encourage the mice to lick and to remain motivated. The first day consisted of 150 trials and subsequent days of 250. The reward, as in the TPSD main task, was dispensed at 1.2 s and remained available to the mice until 2 s, at which time it was sucked away by a vacuum. The mice were required to learn to associate a water reward soon after the stimulus was presented and that there was no water reward in the inter-trial interval (4 s period between trials). Initially, during pre-trials, the experimenter pipetted small drops of water onto the lickport to coax the mice to lick. Once the mice learned this and licked with 80% efficiency, they were advanced to the go/no-go task.

**Figure 1:**
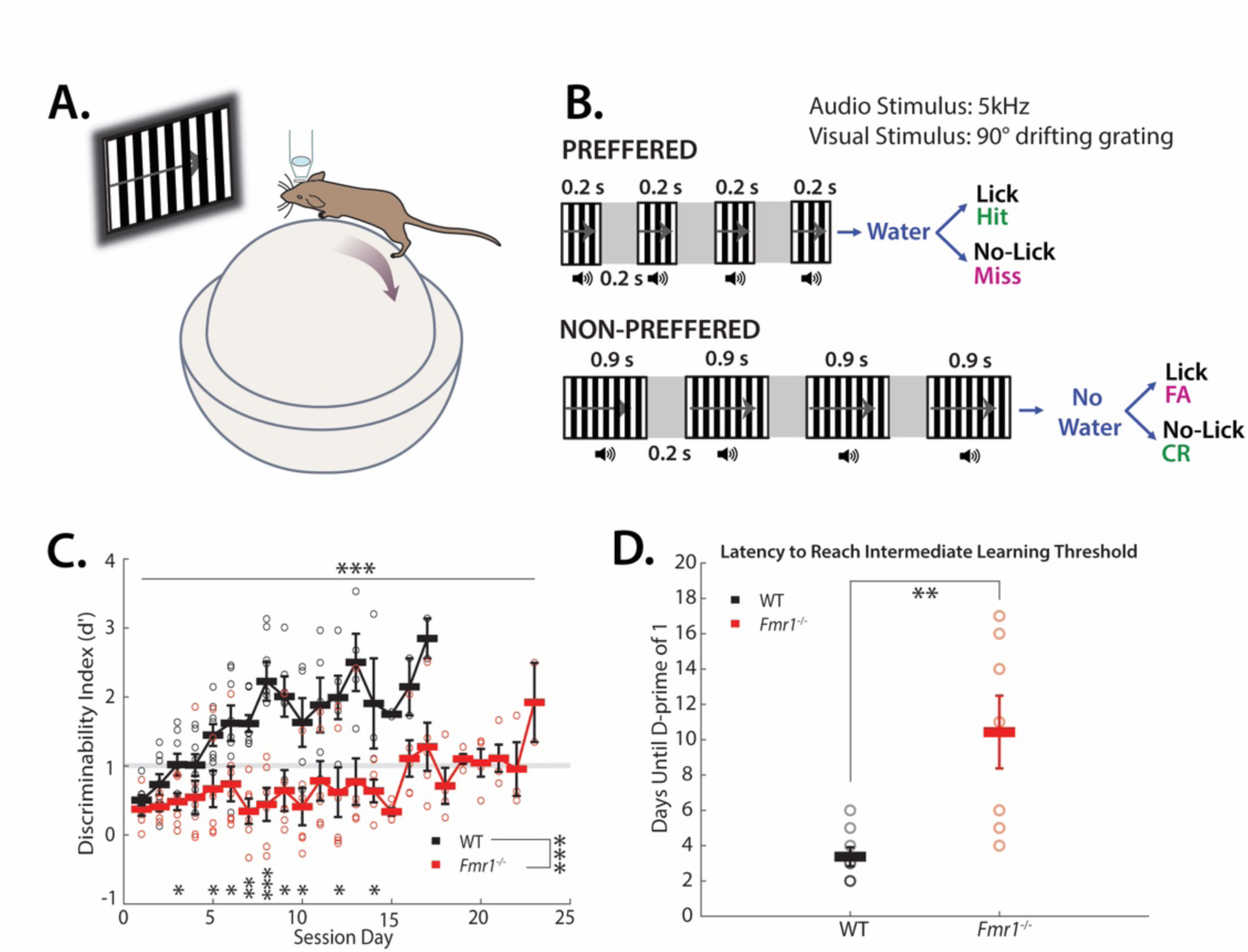
*Fmr1^-/-^* mice exhibit significant learning delays in a timing task (WT n=8; *Fmr1^-/-^*n=7). **A**. Schematic of mouse on polystyrene ball. **B.** Experimental paradigm is a go/no-go task composed of synchronous audiovisual stimuli. **C.** *Fmr1^-/-^* mice exhibited delayed learning of the TPSD (Mixed-effects analysis for training effect: p = 0.0002 and genotype effect: p = 0.0005, followed by Mann-Whitney test for genotype effect at each session: session 1, p = 0.189; session 2, p = 0.108; session 3, p = 0.0426; session 4, p = 0.0721; session 5, p = 0.0401; session 6, p = 0.0205; session 7, p = 0.000666; session 8, p = 0.00470; session 9, p = 0.0303; session 10, p = 0.0303; session 11, p = 0.0823; session 12, p = 0.0190; session 13, p = 0.114; session 14, p = 0.0159; session 15, p = 0.0952; session 16, p = 0.381; session 17, p = 0.0952; Performance is measured by the discriminability index (d’). Grey line at d’ = 1 indicates intermediate learning threshold. **D.** *Fmr1^-/-^* mice take significantly more sessions to reach intermediate learning threshold (d’>1) (on average, 3.38 ± 0.53 d for WT mice vs. 10.43 ± 2.06 d for *Fmr1*^-/-^ mice; p= 0.0037, Student’s t-test)

The TPSD task is a go/no-go task composed of two sequences of synchronous audio- visual stimuli (**Fig. 1B**). Visual stimuli are 90° drifting sinusoidal gratings and are accompanied by a synchronous 5 kHz tone at 80 dB. Within each sequence, four stimuli are presented that differ only in temporality. Our preferred sequence is composed of 4 stimuli of 200 ms; our nonpreferred sequence is composed of 4 stimuli of 900 ms. Each set of the sequences is separated by a 200 ms period of silence accompanied by a grey screen. A water reward is dispensed at 1.2 s and remains available until 2 s, at which time it is sucked away by a vacuum. A custom-built lickport (UCLA engineering) dispensed water, vacuumed it, and recorded licking via breaks in an infrared (IR) beam. Breaks were recorded at 250 Hz. The window in which mice’s licking counts toward a response is 1 to 2 s in both stimuli. A time-out period (6.5 to 8 s), in which the monitor shows a black screen and there is silence, is instituted if the mouse incorrectly responds. The first session was composed of 250 trials and subsequent days of 350. Depending on the stimulus presented, the animal’s behavioral response was characterized as “Hit”, “Miss”, “Correct Rejection” (CR), or “False Alarm” (FA) (**Fig. 1B**). An incorrect response resulted in the time-out period.

To expedite learning, we set the ratio of preferred to nonpreferred stimuli to 70:30 as we found that mice are more prone to licking (providing a ‘yes’ response) than to inhibiting licking (providing a ‘no’ response). We additionally instituted an individualized lick rate threshold to encourage learning as we found that lick rates differed significantly from mouse to mouse. Licking thresholds were calculated from lick rates for mice and showed no significant correlation between licking thresholds and learning rates (WT: Pearson’s *r*, r = .4684, p = -.3012; *Fmr1*^-/-^: Pearson’s *r*, r = - 0.0474, p = 0.9195). This indicates that the individualized lick rate threshold was used as a learning aid to complete the task and did not affect their learning rates or their reliance on the stimulus for task completion. To confirm that mice learned rather than took advantage of the biased 70:30 preferred to nonpreferred trial ratio, we tested mice for 2 additional sessions using a 60:40 ratio of preferred to nonpreferred stimuli. We retain a greater number of preferred stimuli as the total time mice encounter preferred stimuli is less than that of encountering nonpreferred stimuli within a 60:40 trial session (294 s vs 588 s respectively).

On days either side of the 60:40 trial session, *Fmr1^-/-^*mice performed two alternate unimodal sessions, the order of which was randomly shuffled, in which either the visual grating or auditory tone was presented in isolation, and performance was assessed (**Fig. 4A**). WT mice performed a single unimodal session preceding the 60:40 trial session. Following, mice performed a control task, during which visual and auditory stimuli were not presented. Our data shows that mice did not show a learned performance, indicating that they relied on the sensory stimuli for task completion.

Custom-written routines and Psychtoolbox in MATLAB were used to present the visual stimuli, to trigger the lickport to dispense and retract water, and to acquire data.

### Data analysis

#### Discriminability index (d’) and CR and Hit Rates

d’ was calculated using the MATLAB function *norminv*, which returns the inverse of the normal cumulative distribution function: 

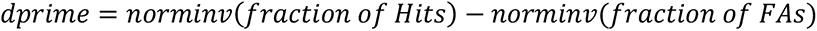

If either rate reached 100% or 0%, we arbitrarily changed the value to either 99% or 1%, respectively. We did this to avoid generating z-scores of infinity that would inaccurately characterize the mice’s performance.

The *d’* of the best 150 trials were selected by a sliding 150 trial window; the highest value was then selected. CR and Hit rates use the same best 150 trial interval.

#### Licking Thresholds

Licking thresholds for each mouse were determined by using the average licking in the last Pretrial session minus 1 standard deviation.

#### Licking Probabilities

Probabilities were taken by binning licks per 0.1s window per trial per mouse. We then averaged each mouse’s probability per time to generate a distribution of probabilities based on trial session, stimulus type, and trial outcome. We use the best 150 trials from each day and each mouse as determined by the discriminability index (*d’*).

#### Support Vector Machine (SVM)

We used the SVM available in the MATLAB Machine Learning and Deep Learning toolbox via the function *fitcsvm*. We used a radial basis function as the kernel. 80% of our data was applied to training the machine and 20% applied to testing it. Instead of training one machine, we developed a strategy wherein we performed a bootstrapped SVM per time per mouse. This allowed us to generate a distribution of accuracy percentages per time to locate critical times of difference during stimulus presentation. 10000 machines were generated per time per mouse for the licking predictor and then averaged as one grand distribution. The licking predictor consisted of binning licks per 0.067 s window per trial per mouse with either the stimulus type (preferred or nonpreferred) or trial outcome (Hit, Miss, CR, FA) as the outcome. With our licking data, we performed no pretraining optimization as we essentially were testing the accuracy of individual features, i.e. time bins of licking.

### Statistical analyses

Statistical analysis of normality (Lilliefors) were performed on each data set and depending on whether the data significantly deviated from normality (p<0.05) or did not deviate from normality (p>0.05) appropriate non-parametric or parametric tests were performed. The statistical tests performed are mentioned in the text and the legends. For parametric two-group analyses, a Student t-test (paired or unpaired) was used; for parametric multi-group analyses, a one-way ANOVA was used. For non-parametric tests, we used the following: Wilcoxon rank sum test (two groups), Kolmogorov-Smirnov test (two groups), and Kruskal-Wallis test (multi-group). When multiple two-group tests were performed, a Bonferroni Correction was applied to readjust the Alpha value. In the figures, significance levels are represented with the following convention: * for p < Alpha; ** for p < Alpha/10, *** for p < Alpha/100. Alpha values are .05 unless otherwise specified. In all the figures, we plot the standard error of the mean (s.e.m.). Graphs either show individual data points from each animal or group means (averaged over different mice) superimposed on individual data points.

For data where we plot the licking profiles in **Figs. 2-3**, we used confidence intervals to significance tests as our data were time series. In addition, recent research has begun to shift away from null-hypothesis significance testing to methods like effect sizes and confidence intervals (Cumming, 2014).

**Figure 2:**
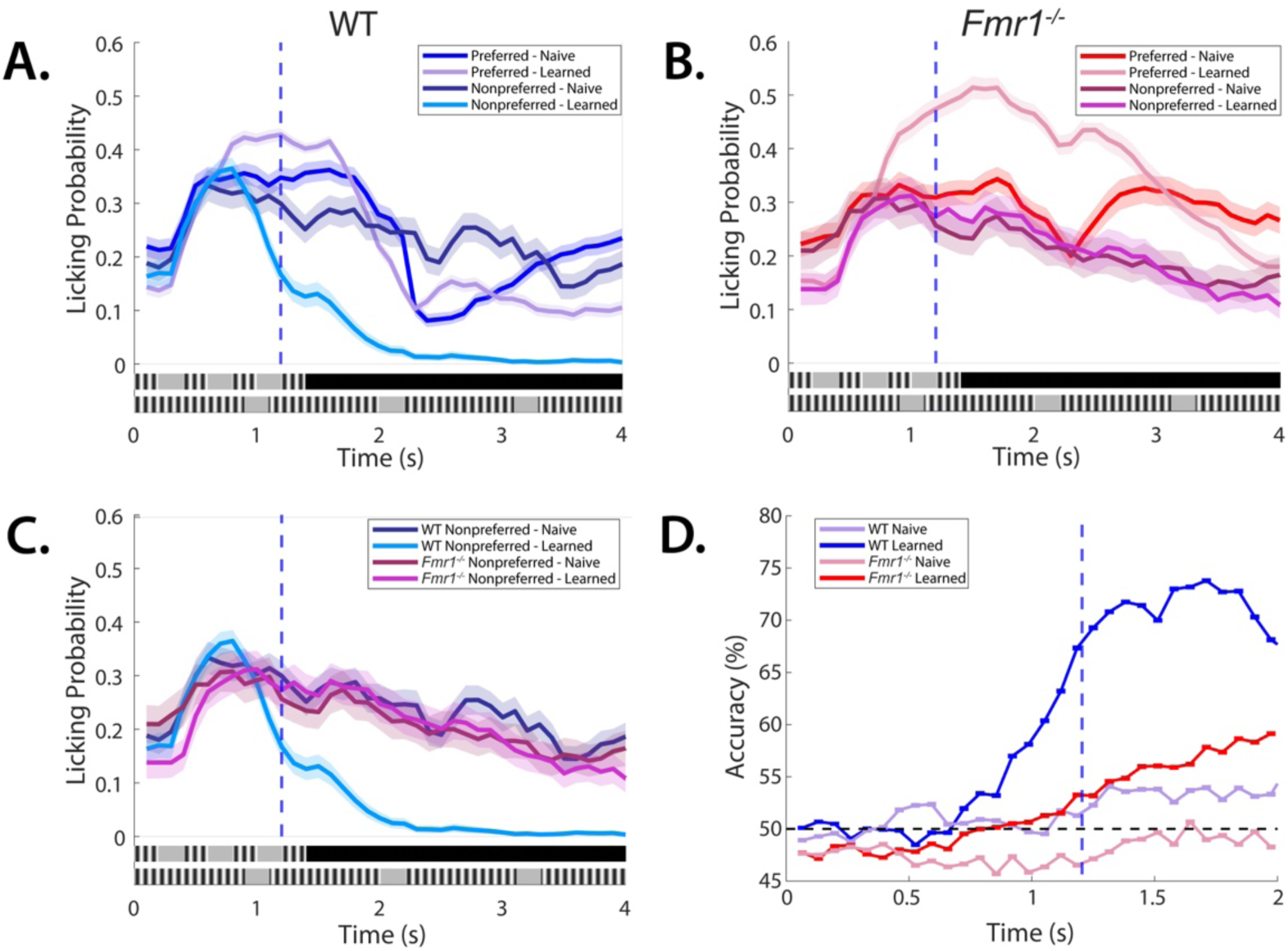
*Fmr1^-/-^* mice demonstrate atypical licking, characteristic of diminished stimulus discrimination and behavioral inhibition. **A.** Probability of a lick event by stimulus type and session for WT mice, shaded areas represent 95% confidence intervals, blue dashed line indicates time of water reward. **B.** Probability of a lick event by stimulus type and session for *Fmr1^-/-^*mice, shaded areas represent 95% confidence intervals, blue dashed line indicates time of water reward. **C.** Probability of a lick event during Non-preferred stimulus by session for both *Fmr1^-/-^* and WT mice, illustrating the absence of refinement in licking response for *Fmr1^-/-^* mice across learning, shaded areas represent 95% confidence intervals, blue dashed line indicates time of water reward. **D.** Accuracy of bootstrapped SVM as a function of time for both *Fmr1^-/-^* and WT mice. Licking events per 0.067 s are the predictors, and stimulus type is the outcome. Learned session accuracy confirms learning as predictability rises above chance before the water reward at 1.2 s (blue dashed line) for WT mice, and confirms diminished learning in *Fmr1^-/-^* mice with predictability being significantly lower at the same timepoint.

**Figure 3:**
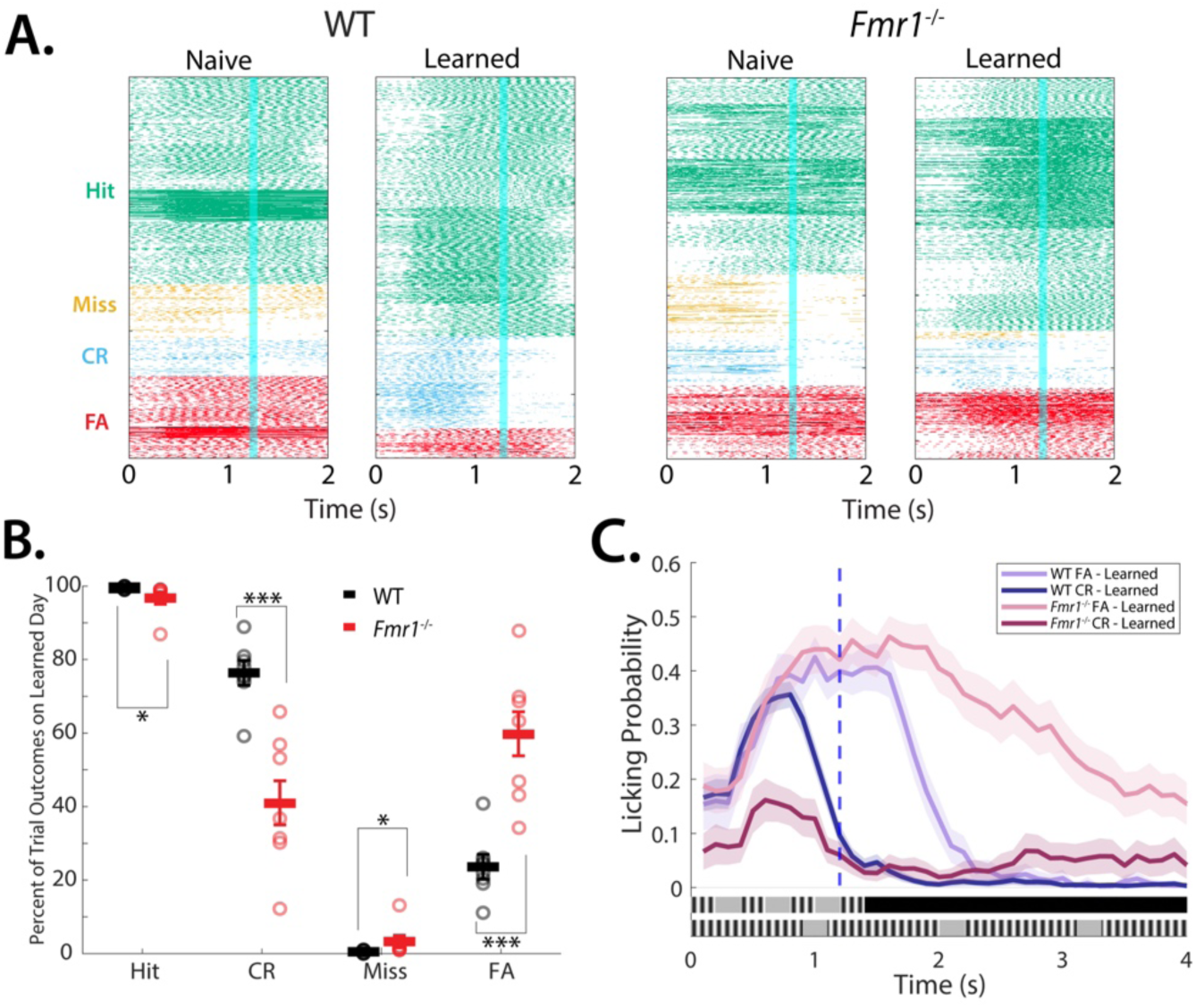
*Fmr1^-/-^*mice demonstrate reduced ability to withhold licking. **A.**, Raster plots of licking in naive and learned sessions in both *Fmr1^-/-^*and WT mice, broken up by trial outcome. Blue line indicates time of water reward. **B.** Percent of trial outcomes for each stimulus type (preferred: Hit/Miss; non-preferred: FA/CR) on expert/best performing session for both *Fmr1^-/-^*and WT mice (on average, FA rate: 23.60 ± 2.97 % for WT mice vs. 59.10 ± 7.02 % for *Fmr1*^-/-^ mice; p = 6.22 x 10^-4^, Mann-Whitney test; CR rate: 76.40 ± 2.97 % for WT mice vs. 40.90 ± 7.02 % for *Fmr1*^-/-^ mice; p = 6.22 x 10^-4^, Mann-Whitney test; Hit rate: 99.50 ± 0.19 % for WT mice vs 96.74 ± 1.67 % for *Fmr1*^-/-^ mice; p = 0.012, Mann-Whitney test; Miss rate: 0.50 ± 0.19 % for WT mice vs 3.26 ± 1.67 % for *Fmr1*^-/-^ mice; p = 0.012, Mann-Whitney test;) **C.** Probability of a lick event during Non- preferred stimulus broken up by trial outcome on expert/best performing session for both *Fmr1^-/-^* and WT mice, shaded areas represent 95% confidence intervals, blue dashed line indicates time of water reward.

**Figure 4:**
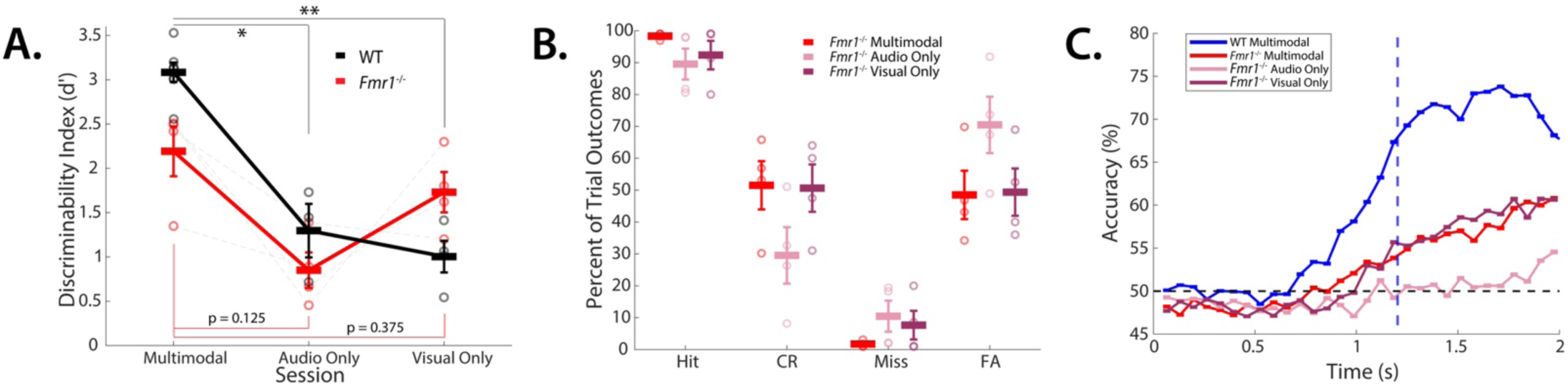
*Fmr1^-/-^* mice display impaired multisensory integration, relying heavily on a single modality. **A.** Performance of both *Fmr1^-/-^* mice and WT mice on multimodal (WT n=7; *Fmr1^-/-^* n=4), compared with audio-only (WT n=3; *Fmr1^-/-^* n=4)(On Average for WT mice d-prime = 3.08 ± 0.110 for multimodal session vs. 1.30 ± 0.302 for auditory-only session; p = 0.0167, Mann-Whitney Test; On Average for *Fmr1*^-/-^ mice d-prime = 2.19 ± 0.282 for multimodal session vs. 0.849 ± 0.201 for auditory-only session; p = 0.125, Wilcoxon signed-rank test), and visual-only sessions (WT n=4; *Fmr1^-/-^* n=4)(On Average for WT mice d-prime = 3.08 ± 0.110 for multimodal session vs. 1.00 ± 0.179; p = 0.0061 for visual-only session, Mann-Whitney Test; On Average for *Fmr1*^-/-^ mice d-prime = 2.19 ± 0.282 for multimodal session vs. 1.73 ± 0.228 for auditory-only session; p = 0.375, Wilcoxon signed-rank test). **B.** Percent of trial outcomes for each stimulus type (preferred: Hit/Miss; non-preferred: FA/CR) on multimodal, audio- only, and visual-only sessions for *Fmr1^-/-^* mice. **C.** Accuracy of bootstrapped SVM as a function of time for both *Fmr1^-/-^* mice across multimodal, audio-only, and visual-only sessions, compared with WT expert multimodal sessions. Licking events per 0.067 s are the predictors, and stimulus type is the outcome. Significant overlap between *Fmr1^-/-^*multimodal and visual-only around time of water reward (blue dashed line) indicates similar levels of stimulus discrimination, while audio-only remains around chance indicating a lack of stimulus discrimination found in other two conditions.

### Exclusion of mice

5 WT and 3 *Fmr1*^-/-^ mice were excluded from the data because the mice lost > 25% body weight (a criterion we established a priori). This had adverse effects on their health that manifested in listlessness, reduced grooming and interaction with cage mates, and occasionally, seizures. One additional *Fmr1*^-/-^ mouse was excluded due to failure to exceed the intermediate learning threshold (D-prime > 1) within three weeks of main-task training.

## Data availability

All the analyzed data reported in this study is available from the corresponding author upon request.

## Code availability

All code including SVM analysis used in this manuscript is available from the corresponding author upon request.

## Results

### *Fmr1*^-/-^ mice exhibit learning impairments on the multimodal temporal pattern sensory discrimination task

To examine temporal pattern learning we have designed a novel go/no-go, Temporal Pattern Sensory Discrimination (TPSD) task (see methods) (Post et al., 2023). Awake, head-restrained young adult mice (2-3 months) are habituated to run on a polystyrene ball treadmill while they perform the TPSD paradigm. Water-deprived mice are presented with two audio-visual temporal patterns (preferred and non-preferred) as shown in the schematic in **Fig. 1B**. Each pattern consists of 4 audio-visual (AV) stimuli, where each AV stimulus lasts either 0.2 s or 0.9 s and is separated by a 0.2 s gray screen. The visual stimulus consists of a drifting sinusoidal 90° grating, and the auditory stimulus consists of a 5kHz tone. Both auditory and visual stimuli are presented concurrently; therefore, the stimuli are audio-visual. The temporal pattern with 0.2 s AV stimuli is associated with a water reward (preferred pattern) and the temporal pattern with 0.9 s AV stimuli is not (nonpreferred pattern) (**Fig. 1B**). We quantified the performance of mice using a discriminability index (*d’*) in which d’ = 1 was set as an intermediate learning threshold (**Fig. 1C, D**). Compared to wild-type (WT) control mice, *Fmr1*^-/-^ mice demonstrated marked impairment in task performance. First, they exhibited a delay in reaching the intermediate learning threshold (**Fig. 1C**: Mixed effects analysis for training effect: p = 0.0002 and genotype effect: p = 0.0005, followed by Mann-Whitney test for genotype effect at each session; **Fig 1D**: on average, 3.38 ± 0.53 d for WT mice vs. 10.43 ± 2.06 d for *Fmr1*^-/-^ mice; p= 0.0037, Student’s t-test). Second, we quantified learning using a d’ = 2, indicative of expert performance, as have previously done (Goel et al., 2018; Post et al., 2023; Rahmatullah et al., 2023). Wild-type (WT) mice achieved a d’=2 by preferentially licking to the preferred pattern and withholding licking for the non-preferred pattern. As we have previously observed with *Fmr1*^-/-^ mice, the learning capacities of the mice differed. Compared to WT mice, we found a bimodal distribution in the expert performance of *Fmr1*^-/-^ mice (**Supp.** Fig. 1A). While one group of *Fmr1*^-/-^ mice failed to reach expert status even after two weeks of additional sessions of training beyond the WT mean, another group achieved expert status with an average delay of an additional week of training (on average, 8.38 ± 1.35 d for WT mice vs. 15.0 ± 3.63 d for *Fmr1*^-/-^ mice; p= 0.0598, Student’s t-test**; Supp.** Fig. 1B). Interestingly, 2 of these 4 *Fmr1*^-/-^ mice that achieved expert status (d’>2), did so within the same timeline as the WT mice, potentially indicative of a high-functioning phenotype.

Prior to training on the TPSD task, mice perform the pretrial task. During the pretrial task mice experience only the preferred stimulus and every trial is rewarded. This allows mice to learn to lick reliably (>80% licking) and learn the task structure–association of stimulus with water reward (see Methods). This pretrial task is similar to previous studies (Goel et al., 2018; Post et al., 2023; Rahmatullah et al., 2023) and is a common strategy used in behavior assays(Guo et al., 2014). While *Fmr1*^-/-^ mice showed impairments in learning the TPSD task, there was no difference in their ability to perform the pretrial task (on average, 5 ± 0.71 d for WT mice vs. 6.29 ± 1.36 d for *Fmr1*^-/-^ mice; p= 0.398, Student’s t-test, **Supp.** Fig. 2). Interestingly, both genotypes exhibited a positive *d’* of 0.5 on session 1 of the TPSD task, which likely resulted from mice learning to associate stimulus with reward in the pretrial task prior to the TPSD task. Thus, *Fmr1*^-/-^ mice did not exhibit any defects in learning the task structure, rather the deficits were specific to stimulus discrimination.

### Atypical licking patterns

As mice train on the TPSD task, learning is manifested in the dynamic profile of the licking patterns. Both genotypes showed no change in licking profile between preferred and non-preferred stimuli, on the naive session (**Figs. 2A-B**). WT mice demonstrated a sustained increase in licking to the preferred stimulus, with a concomitant decrease in licking to the non-preferred stimulus (**Fig. 2A**). *Fmr1*^-/-^ mice revealed a sustained elevation in licking to the preferred stimulus, however, licking to the non-preferred stimulus was also prolonged, and lasted well into the trial (**Fig. 2B**). The difference in licking profiles of the WT and *Fmr1*^-/-^ mice to the non-preferred stimulus is underscored in **Fig. 2C**.

### Lack of predictability in licking profiles from *Fmr1*^-/-^ mice

We have previously developed a bootstrapped Support Vector Machine (SVM), a type of binary classifier to establish causality between licking profiles and 1) performance; and 2) demonstrate that the dynamic nature of licking profiles accompanies changes in learning (Post et al., 2023; Rahmatullah et al., 2023). Using a similar classifier, we compared the accuracy of licking in *Fmr1*^-/-^ with WT mice. We run our SVM 10,000 times within 0.067 s time bins using licking within a trial as the predictor and the trial stimulus type as the outcome (see Methods). This allows us to generate a distribution of correctly predicted outcomes per time bin per mouse, which is then compared to a randomly shuffled control (see Methods). **Fig. 2D** shows that in both genotypes, on naïve sessions, the predictability of licking is at chance. There is a small increase in predictability following the delivery of the water reward. Licking profiles in expert WT mice show a robust increase in predictability of preferred stimulus, well before delivery of water.

This suggests that learned mice expect to receive a water reward on trials with preferred stimuli. Predictability of licking patterns in *Fmr1*^-/-^ mice continues to hover over chance indicating that licking responses in *Fmr1*^-/-^ mice are not related to stimulus type. This further exacerbates the diminished learning displayed by *Fmr1*^-/-^ mice (as measured by d’ values).

### Reduced CR rates as a measure of impaired learning

d’ metric used to gauge performance is a ratio of Hit rate and False Alarm (FA) rate. After competition of the pretrial task, when mice begin training on the TPSD task, they achieve high Hit rates, since they have been trained to lick to the preferred stimulus in the pretrial task. On the TPSD task, an increase in d’ is achieved by lowering the FA rate (or increasing the CR rate). The ratio of preferred to nonpreferred stimulus ratio can artificially amplify the effect of the Hit rate on mice’s *d’* value. Therefore, we also calculated the changes in Hit, CR, and FA rates. Licking raster plots obtained from all the mice show no change in CR and FA rates with task performance in *Fmr1*^-/-^ mice (**Fig. 3A**). Thus, compared to WT mice, *Fmr1*^-/-^ mice showed a significant increase in FA rates, and coordinated decrease in CR rates, as well as a decreased Hit-rate and increased Miss rate ( on average, FA rate: 23.60 ± 2.97 % for WT mice vs. 59.10 ± 7.02 % for *Fmr1*^-/-^ mice; p = 6.22 x 10^-4^, Mann-Whitney test; CR rate: 76.40 ± 2.97 % for WT mice vs. 40.90 ± 7.02 % for *Fmr1*^-/-^ mice; p = 6.22 x 10^-4^, Mann-Whitney test; Hit rate: 99.50 ± 0.19 % for WT mice vs 96.74 ± 1.67 % for *Fmr1*^-/-^ mice; p = 0.012, Mann-Whitney test; Miss rate: 0.50 ± 0.19 % for WT mice vs 3.26 ± 1.67 % for *Fmr1*^-/-^ mice; p = 0.012, Mann-Whitney test; **Fig. 3B**.). Since several *Fmr1*^-/-^ mice did not achieve expert performance on the TPSD task, learned day for the *Fmr1*^-/-^ was the final session that they were trained on. We further examined the licking profiles specifically on FA and CR trial outcomes. WT mice displayed an increase in licking in both Hit and FA trials, however, the licking was sharply reduced in CR trials. Compared to WT mice, *Fmr1*^-/-^ mice displayed prolonged licking in FA trials. An overall reduction in the licking magnitude of *Fmr1*^-/-^ mice on CR trials, paired with significantly lower CR rates compared to WT controls, is a further indication of the difficulty of withholding licking in this model, suggestive of hyperarousal phenotype (Goel et al., 2018; Rahmatullah et al., 2023).

### Multimodal nature of stimulus does not aid learning in *Fmr1*^-/-^ mice

Neurotypical humans and rodents perform better with multimodal stimuli than with unimodal stimuli (Raposo et al., 2012; Barakat et al., 2015). However, our data showed an impairment in TPSD where multimodal stimuli were utilized. Further, impairment in audio-visual integrations has been observed in ASD(Foss-Feig et al., 2010; Stevenson et al., 2014a; Stevenson et al., 2014b). Therefore, we reasoned that unimodal stimuli might improve the performance of *Fmr1*^-/-^ mice. Therefore, after mice were trained on the multimodal TPSD task, they were then tested on 1 session of the task where the stimulus presented was unimodal–either auditory only or visual only. Across both genotypes, we observed a marked decrease in performance when the stimuli consisted of auditory stimuli only (Average d-prime for WT mice = 3.08 ± 0.110 for multimodal session vs. 1.30 ± 0.302 for auditory-only session; p = 0.0167, Mann-Whitney Test; Average d-prime for *Fmr1*^-/-^ mice = 2.19 ± 0.282 for multimodal session vs. 0.849 ± 0.201 for auditory-only session; p = 0.125, Wilcoxon signed-rank test). *Fmr1*^-/-^ mice show a trend for increased FA rate and reduced CR rate, in agreement with the reduced d-prime on the auditory only version of the task (**Fig. 4B**). However, when *Fmr1*^-/-^ mice performed the TPSD task consisting of visual-only stimuli, there was very little difference in performance between the multimodal and visual-only versions of the task, whereas WT mice performed at a significantly lower level compared to their multimodal task level (Average d-prime for WT mice = 3.08 ± 0.110 for multimodal session vs. 1.00 ± 0.179; p = 0.0061 for visual-only session, Mann-Whitney Test; Average d-prime for *Fmr1*^-/-^ mice = 2.19 ± 0.282 for multimodal session vs. 1.73 ± 0.228 for auditory-only session; p = 0.375, Wilcoxon signed-rank test). This is especially apparent when looking at SVM stimulus predictivity from licking (**Fig. 4C**), wherein the predictive accuracy for the *Fmr1*^-/-^ mice is nearly inseparable just preceding the water reward across multimodal and visual-only sessions, while predictivity in the auditory-only sessions remains at or around chance. This suggests that *Fmr1*^-/-^ mice relied predominantly on the visual stimulus and the presence of the synchronous auditory tone in the multimodal stimuli did not aid in stimulus discrimination.

## Discussion

Distortions in timing and time perception are often observed in FXS and ASD, however, the neural basis of the timing difficulties is underexplored (Allman and DeLeon, 2009; Allman and Meck, 2012; Hinault et al., 2023). Here we tested the performance of a mouse model of FXS (*Fmr1*^-/-^ mouse) on a novel timing task–TPSD, where mice have to discriminate between durations of audio-visual stimuli to reach expert status (Post et al., 2023). Our results show severe impairments in performance on the TPSD task, where mice exhibited a delay in, or an inability to reach expert status. Task performance is measured by preferential changes in licking to the preferred stimulus duration. Using multiple analyses, we found that *Fmr1*^-/-^ mice revealed an impairment in licking profiles, indicative of an inability to discriminate between durations of time. Interestingly, while WT mice benefited from the multimodal nature of stimuli in the TPSD task, trends in our data show that *Fmr1*^-/-^ mice rely on visual stimuli alone. This suggests a disruption in multisensory integration which has been reported in the context of FXS and ASD (Stevenson et al., 2014b; Wallace and Stevenson, 2014). Together our results establish a timing task that allows investigation of timing difficulties in a mouse model.

### Impaired time perception and temporal processing in FXS and related neurodevelopmental disorders

Impaired sensory processing is a core symptom of ASD and FXS and the sensory atypicalities range from auditory hypersensitivity, tactile defensiveness, and disruption in discrimination and perception of sensory stimuli. Difficulties in processing temporal features in sensory stimuli and perceiving time are often observed in ASD, Attention Deficit Hyperactivity Disorder (ADHD), and FXS. Research in humans examining temporal processing classifies temporal judgments into four categories: a) discrimination between durations, b) temporal order of durations, c) time perception (how much time has elapsed), and d) mental time travel (Hinault et al., 2023). Difficulties in all these categories have been reported in ADHD, FXS, and ASD. Particularly, subjects with ADHD overestimate time, judging durations to be longer than Typically developing Controls (TDC). Other studies show that ADHD individuals underproduce durations of time, judging durations to be shorter (Walg et al., 2017; Zheng et al., 2022). Timing impairments have also been identified in FXS individuals ranging from infants and toddlers to young adults (Kogan et al., 2004; Hall et al., 2009; Farzin et al., 2011). Hall et al, 2009 investigated the underlying neural dysfunction that accompanied timing deficits during an auditory discrimination task and identified increased activation of neural pathways in the left hemisphere and superior temporal gyrus in girls with FXS. More recently studies using *Fmr1*^-/-^ mice have shown dysfunctional temporal processing specific to earlier stages of development with a higher prevalence in female mice (Croom et al., 2023; Croom et al., 2024).

### Temporal processing and speech

Delayed language development is a core symptom of ASD (Abbeduto and Hagerman, 1997; Abbeduto et al., 2007). Given that temporal processing is critical for speech and language function (Shannon et al., 1995), meaning in spoken language, and production of speech derives from sequences of syllables that are highly temporally structured, two important variables need to be solved: 1) Understanding the defects in temporal processing in FXS and ASD. 2) How information about time is extracted from sensory stimuli and encoded in neural dynamics is important in solving deficits in language development. A recent study examined the neural dynamics associated with impaired temporal processing in *Fmr1*^-/-^ mice, using a gap-ASSR paradigm (Rumschlag and Razak, 2021; Croom et al., 2024). This paradigm determines the cortex’s ability to phase lock to gaps in auditory stimuli, thus providing a function of temporal processing. Croom et al, 2024 found that female *Fmr1*^-/-^ mice exhibited earlier maturation of temporal processing and an elevation in ERP amplitudes through development. Our novel TPSD paradigm addresses the second important unknown, where we investigate timing impairments in the context of a task– mice have to learn to discriminate between durations of time to receive a water reward. Thus, mice must learn to extract temporal structure from the stimuli to achieve expert status. We recently demonstrated, in WT mice, the emergence of specific neural trajectories that contain information about the temporal structure of stimuli (Post et al., 2023). Disruptions in selectivity and modulation of primary visual cortical (V1) cells have been shown to be associated with impaired sensory learning in *Fmr1 ^-/-^* mice (Goel et al., 2018; Rahmatullah et al., 2023). We expect similar impairments in V1 neural dynamics in *Fmr1^-/-^* mice, where pyramidal cells will fail to develop distinct activity patterns through learning on the TPSD task.

Recent work suggests that speech deficits are associated with a lack of coherence in neural activity between auditory cortical areas and frontal cortex (Schmitt et al., 2020). Whether speech deficits result from reduced inter-cortical communication or is a combination of dysfunction in local and long-range communication remains to be determined. Further, speech as a form of communication is strongly associated with social cues (Baldwin and Tomasello, 1998; Lee and Lew-Williams, 2022). Difficulties in social cognition are another hallmark of FXS and ASD and indeed have been investigated in multiple mouse models. Difficulties in language development can result in social deficits, however, it is unknown if a common neural dysfunction is associated with atypical behaviors.

### Multisensory integration

Sensory stimuli are often multimodal, there are sensory receptors and neural mechanisms in place to extract information from each modality. Indeed, typically developing (TD) humans benefit from combining multimodal stimuli for task performance (Alais et al., 2010) and this has been shown in rodent studies too (Raposo et al., 2012). Individuals with ASD do not exhibit a similar benefit in task performance with multisensory stimuli, likely due to disruption in multisensory integration (Smith and Bennetto, 2007; Irwin et al., 2011; Bebko et al., 2014; Stevenson et al., 2014a; Stevenson et al., 2014b; Foxe et al., 2015). While some studies suggest that deficit in multisensory integration is restricted to early development, one study showed that teenagers with ASD showed reduced benefits from audio-visual stimuli (Brandwein et al., 2013). Further, in a visual search task, individuals with ASD showed increased performance in the absence of an auditory tone that was designed to facilitate task performance (Collignon et al., 2013). In contrast, TD controls showed a gain in performance in the presence of the auditory tone. In our study, *Fmr1*^-/-^ mice exhibited a similar phenotype resulting in a trend for a diminished reliance on paired audio-visual stimuli in the TPSD task. Further, the performance of *Fmr1*^-/-^ mice on the unimodal task suggested that auditory stimuli tended to impede rather than facilitate task performance. One hypothesis is that simple audio-visual stimuli, similar to the stimuli in the TPSD paradigm are encoded in cross-modal interactions across primary auditory and visual cortices (Molholm et al., 2002). Indeed, several studies in WT mice have also shown that audio-visual stimuli evoke multimodal plasticity in primary visual cortex (V1) (Morrell, 1972; Petro et al., 2017; Garner and Keller, 2022). Future experiments will determine whether deficits in plasticity mechanisms across sensory cortices impair the encoding of audio-visual stimuli.

Focusing basic and translational research efforts in animal models of FXS can help identify dysfunctional neural mechanisms, contributing to deficits. However, designing rodent behaviors, such as the TPSD assay, to capture sensory symptoms observed in humans will be essential for developing effective interventions for FXS and ASD.

## Author contributions

W.M. and A.G. conceived the project and designed the experiments.

W.M. and S.P. wrote the MATLAB code for data acquisition and analysis with help from A.G.

W.M. conducted the experiments and analyzed the data. A.G. and W.M. interpreted the data and wrote the paper with input from other authors.

## Acknowledgments

The authors thank Bart Kats for help with using the Nautilus clusters for running the SVM analysis.

