## Supplementary Figures for "Learning Impairments in *Fmr1^-/-^* mice on an Audio-Visual Temporal Pattern Discrimination Task"

**Supplemental Material**

**Supplementary Figure 1:** *Fmr1^-/-^* mice show variability in learning capacity, splitting into two groups of those who achieve expert performance (n = 4) and those who fail to do so (n = 3). **A.** Scatter of performance across all main trial sessions, color-matched per individual mouse, with mean. Performance is measured by the discriminability index (d’). **B.** Number of sessions taken to reach expert performance (d’>2) for *Fmr1^-/-^* (n = 4) and WT mice (n = 8)(on average, 8.38 ± 1.35 d for WT mice vs. 15.0 ± 3.63 d for *Fmr1*^-/-^ mice; p= 0.0598, Student’s t-test).

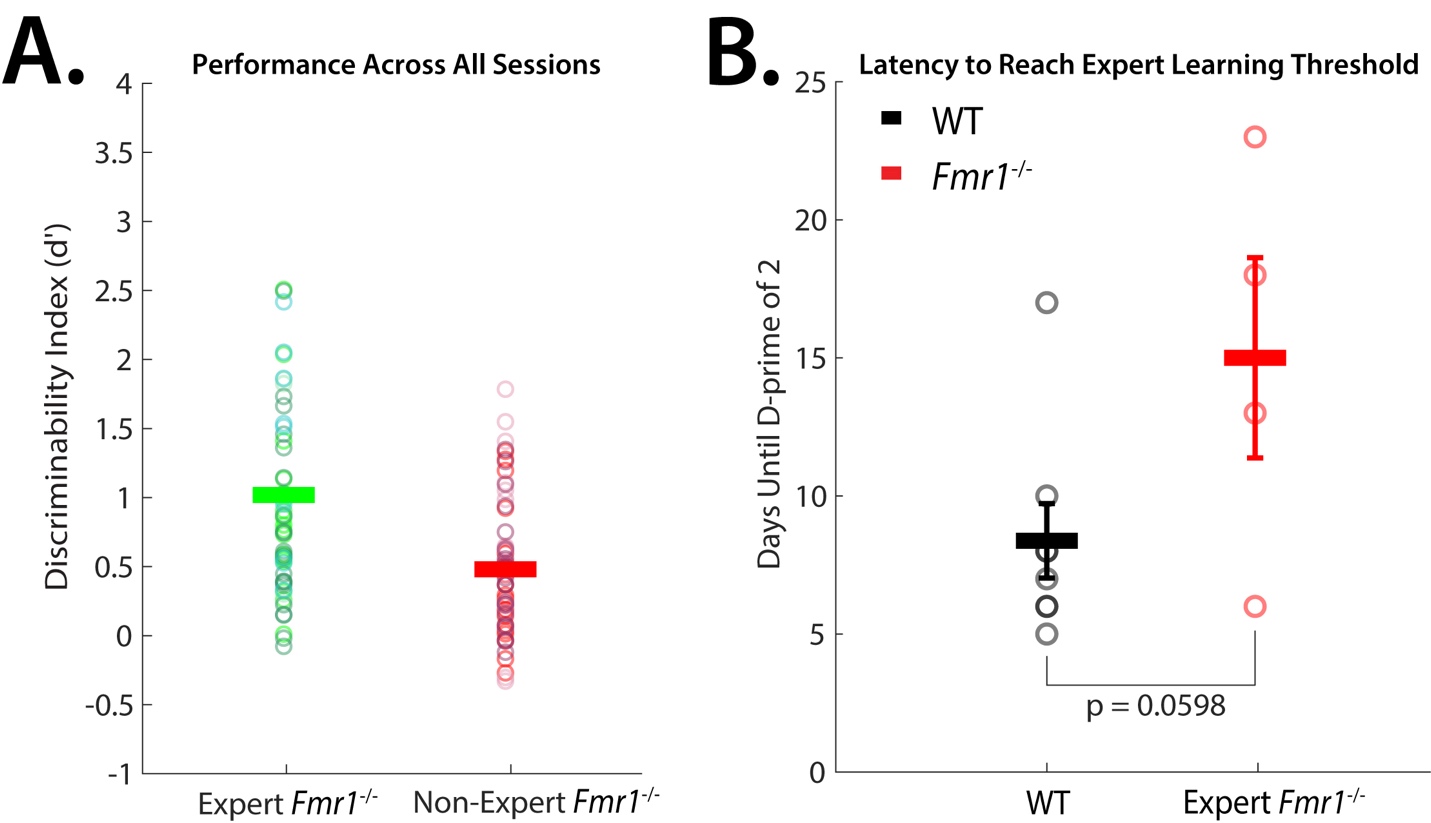

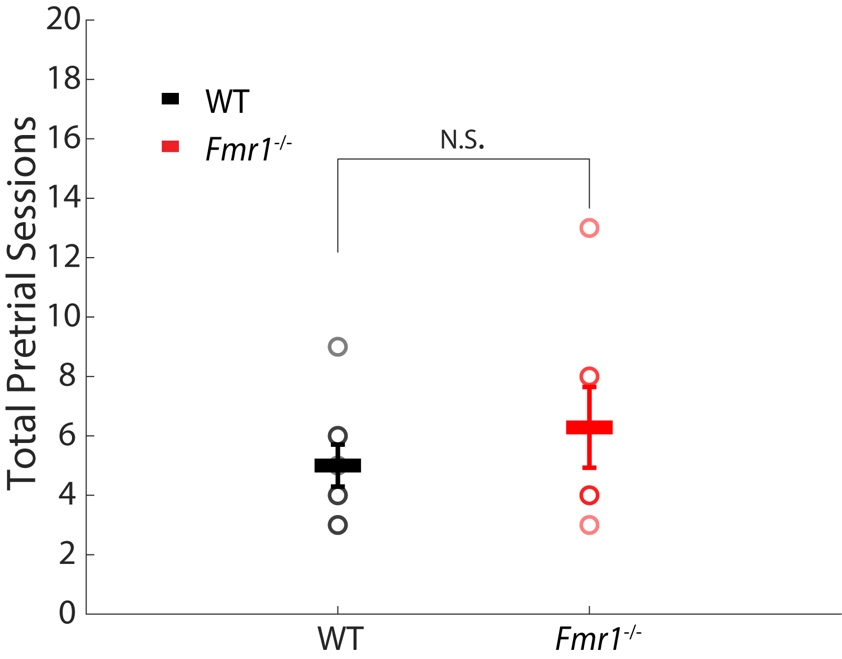

**Supplementary Figure 2:** WT and *Fmr1^-/-^* mice show no differences in pretrial performance. Number of sessions taken to progress past pretrials (on average, 5 ± 0.71 d for WT mice vs. 6.29 ± 1.36 d for *Fmr1*^-/-^ mice; p= 0.398, Student’s t-test).
